# An in vitro platform for quantifying cell cycle phase lengths in primary human intestinal stem cells

**DOI:** 10.1101/2023.10.09.561410

**Authors:** Michael J Cotton, Pablo Ariel, Kaiwen Chen, Vanessa A Walcott, Michelle Dixit, Keith A Breau, Caroline M Hinesley, Kasia Kedziora, Cynthia Y Tang, Anna Zheng, Scott T Magness, Joseph Burclaff

## Abstract

**Background and Aims:** The intestinal epithelium exhibits dynamic control of cell cycle phase lengths, yet no experimental platform exists for directly analyzing cell cycle phases in living human intestinal stem cells (ISCs). Here, we develop primary human ISC lines with two different reporter constructs to provide fluorescent readouts to analyze cell cycle phases in cycling ISCs.

**Methods:** 3D printing was used to construct a collagen press for making chamber slides that support primary human ISC growth and maintenance within the working distance of a confocal microscope objective. The PIP-FUCCI fluorescent cell cycle reporter and a variant with H2A-mScarlet that allows for automated tracking of cell cycle phases (PIP-H2A) were used in human ISCs along with live imaging and EdU pulsing. An analysis pipeline combining free-to-use programs and publicly available code was compiled to analyze live imaging results.

**Results:** Chamber slides with soft collagen pressed to a thickness of 0.3 mm concurrently support ISC cycling and confocal imaging. PIP-FUCCI ISCs were found to be optimal for snapshot analysis wherein all nuclei are assigned to a cell cycle phase from a single image. PIP-H2A ISCs were better suited for live imaging since constant nuclear signal allowed for more automated analysis. CellPose2 and TrackMate were used together to track cycling cells.

**Conclusions:** We present two complete platforms for analyzing cell cycle phases in living primary human ISCs. The PIP-FUCCI construct allows for cell cycle phase assignment from one image of living cells, the PIP-H2A construct allows for semi-automated direct quantification of cell cycle phase lengths in human ISCs using our computational pipeline. These platforms hold great promise for future studies on how pharmaceutical agents affect the intestinal epithelium, how cell cycle is regulated in human ISCs, and more.

## INTRODUCTION

Cells of the intestinal epithelium exhibit dynamic control of cell cycle at homeostasis and following perturbation. Intestinal stem cells (ISCs) and rapidly dividing transit amplifying (TA) cells proliferate at different rates, with their predicted doubling time averaging around 24 hours and 12 hours, respectively (Barker, van Es et al. 2007). Exogenous nutrients and other dietary molecules can alter cell cycle (Calibasi-Kocal, Mashinchian et al. 2021), and cells cycle differently in response to acute or chronic injuries and disease (McKernan and Egan 2015, Rees, Tandun et al. 2020). Determining the effects of these cell cycle changes and how they are regulated is an active field of study. However, no platforms exist in the current literature wherein cell cycle phases can be visualized and tracked in live human ISCs.

Cell cycle has been studied in the intestinal epithelium using multiple techniques, and nearly always using model systems. Early studies tracked proliferative populations of the small intestinal epithelium by quantifying mitotic indices or by pulsing with tritiated thymidine, with studies from the 1970s using per cent labelled mitosis (PLM) or vincristine mitotic accumulation (VCR) experiments in mice and rats to estimate that populations in the small intestinal crypt cycled every 10-24 hours depending on cell position (reviewed in (Potten 1986)). In 2011, Schepers et al. paired EdU pulsing with PH3 immunofluorescent staining to show that mouse ISCs divide every 21.5 hours on average (Schepers, Vries et al. 2011). However, these methods estimate the full cell cycle, without analyzing individual phase lengths. Similarly, studies on cell cycle perturbation often rely on staining for expression of proteins such as Ki67 or PCNA or by pulsing cells with nucleotide analogs such as tritiated thymidine, BrdU, or EdU, which mark cells actively synthesizing DNA at the time of pulsing. While these strategies are useful for comparing overall proliferation between populations, they do not inform about differences in lengths of individual cell cycle phases.

A major current method for analyzing cell cycle phases entails using flow cytometry to analyze DNA content in cells using Hoechst or propidium iodide. Tracking cells with 2n DNA (G1/G0 phase), 4n DNA following full replication (G2/M phase), or an intermediate amount (S phase) can show the proportion of cells in each phase at a given time and serve as a useful comparison between populations. However, this method does not directly measure cell cycle phase lengths and, as such, can miss valuable information. For example, a ratiometric technique such as DNA content analysis would be unable to determine if a certain perturbation shortens or lengthens all phases of the cell cycle – instead it would only inform if one phase is changed more than the others. For more nuanced results like this, or for comparing cell cycle dynamics across individual cells, direct measurement of cell cycle phase lengths holds benefits.

A new strategy for directly measuring cell cycle phase lengths appeared in 2008 with the advent of the FUCCI cell cycle reporter allele (Sakaue-Sawano, Kurokawa et al. 2008). This construct allowed living cells to be delineated into G1 or S/G2/M phases based on expression of fluorescent reporters. Several iterations of this reporter followed, making readouts of phase changes more concise and accurate and allowing for tracking of G1, S, and G2/M phases separately (Zielke and Edgar 2015, Grant, Kedziora et al. 2018). This technique has been used in the intestine, largely utilizing mice engineered to express the reporters across all tissues (Abe, Sakaue-Sawano et al. 2013). Studies have used these powerful reporters to demonstrate that murine ISCs reside largely within the G1 phase (Carroll, Newton et al. 2018) or to show ratios of cells within each cell cycle phase, replicating the results of DNA analysis but in live growing cells (Stokes, Cooke et al. 2017). One study used murine FUCCI2 cells to trace individual cells to track cell cycle length with regards to circadian rhythm (Matsu-Ura, Dovzhenok et al. 2016). We expanded on this model to engineer a platform in which primary human ISCs could be tracked in an easy-to-image monolayer platform.

In this study, we combine use of our primary human ISC monolayer culture system (Wang, DiSalvo et al. 2017, Wang, Gunasekara et al. 2017) with two engineered cell cycle reporters to allow for cell cycle phases to be directly measured in freely cycling ISCs by either snapshot analysis of single images or by live imaging to directly quantify phase lengths. To make this possible, we 1) developed a collagen molding process to optimize chamber slides for confocal microscopy of primary ISCs, 2) generated two genetically engineered primary human ISC cell cycle reporter lines, and 3) developed a computational analysis pipeline using publicly available software. Importantly, by using only two fluorescent channels and utilizing publicly-available analysis software, our platform maintains both high adaptability to a range of projects and an easy-to-implement and open-source analysis pipeline.

## RESULTS

### An engineered press eliminates the collagen meniscus for optimal imaging

Creating a highly reproducible and homogenous ISC culture platform for cell cycle analysis of primary human ISCs requires several design features that would simultaneously permit ISC maintenance and live imaging. Our goal was to minimize the working distance between the microscope objective and the cells to maximize imaging signal and focus while also maintaining a sufficiently thick collagen hydrogel that would create a soft culture substrate (∼9 KPa) necessary to preserve ISC self-renewal and reduce differentiation (Wang, DiSalvo et al. 2017, Wang, Gunasekara et al. 2017). To minimize the working distance for imaging, ISCs were cultured on Ibidi μ-Slide 4 Well chamber slides which have a #1.5 polymer coverslip base (170 μm thick) and four separate chambers to allow for multiple perturbations within the same imaging experiment. When collagen was introduced into the wells, a large meniscus formed creating a curvature that substantially impaired imaging in the z-plane and reduced the thickness of the collagen at the center of the well to the point that the hard polystyrene slide transferred high stiffness through the very thin layer of collagen (Figure 1A). Under this scenario, ISCs near the center of the wells were within the focal range of the objective but the collagen was insufficiently thick to support ISC maintenance and proliferation. This resulted in phenotypic differences in the ISCs, with nuclei spaced further apart and the majority of the cells exiting the cell cycle, as seen by lack of EdU uptake (Figure 1B). By contrast, the edges of the meniscus had thick collagen which supported ISC proliferation, but the cells were not on a flat surface and upper regions of the meniscus were above the focal range of the objective. The differences in proliferation status and cell density across the curvature of the collagen indicated that the prototype system causes heterogenous ISC growth, limiting interpretations of cell cycle states and substantially decreasing reproducibility.

**Figure 1:**
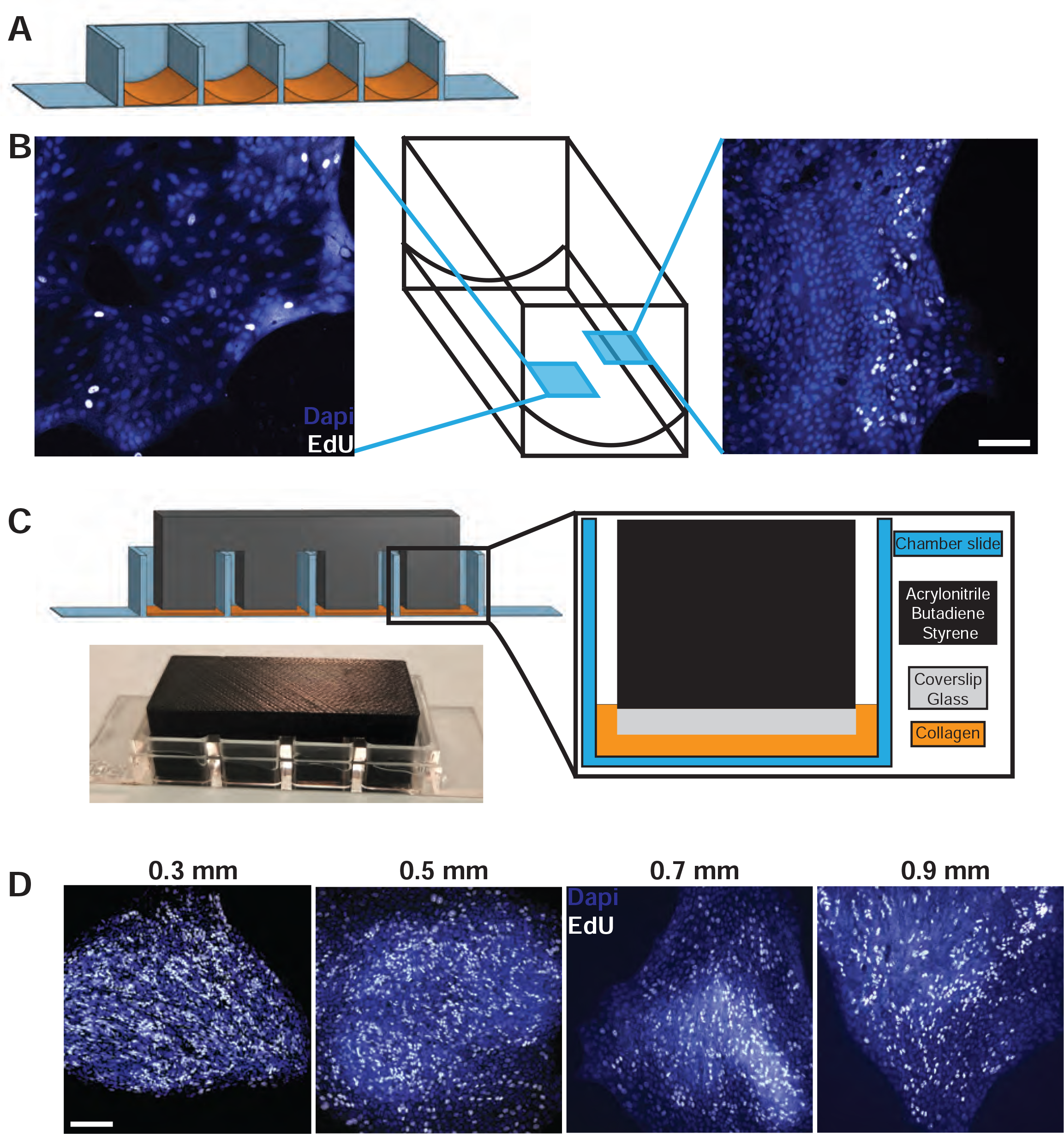
Designing the collagen press. **A)** Model showing meniscus formation following collagen being freely poured into chamber slides than allowed to set. **B)** Immunofluorescence of nuclei (DAPI, blue) and proliferation (EdU uptake, white) in cells at the thin middle or the thick outer edge of the meniscus. Scale bar = 50 μm. **C)** Model for 3D-printed collagen press designed to sit across the chamber slide walls and reach into the wells to prevent meniscus formation. Cover slip glass is glued to the leg bottoms to form a flat surface. Photograph of a collagen press sitting on a chamber slide. **D)** Representative images of cells cultured on collagen patties with thicknesses ranging from 0.3 – 0.9 mm stained for nuclei (DAPI, blue) and proliferation (EdU uptake, white). Scale bar = 100 μm.

To address this issue, we explored methods to create a flat collagen substrate that was sufficiently thin to enable live confocal imaging while supporting ISC self-renewal. We designed a collagen press to hold the surface of the collagen flat while it solidified (Figure 1C). The press was 3D printed (Supplemental File 1), then the surface of each press that contacts the collagen was capped with coverslip-thickness glass to create a flat and even collagen surface and to reduce adhesion to the collagen for easy removal (Figure 1C). The press was designed to rest on the walls separating the slide chambers to create a gap of pre-determined width for the collagen to solidify within. To determine the optimal collagen thickness for imaging, presses with staggered lengths were constructed to test ISC growth on collagen thicknesses ranging from 0.3 mm to 0.9 mm. All four thicknesses supported cell cycle maintenance, as indicated by sustained EdU uptake and closely clustered nuclei after 3 days in culture (Figure 1D). Final presses were designed to form 0.3 mm collagen patties to minimize working distance while maintaining stemness and proliferation. The final platform allowed for maintenance of primary human ISCs on a flat surface with a minimal working distance to allow for use of many confocal microscope objectives, with four wells to allow for multiple concurrent perturbations.

### PIP-FUCCI human ISC reporters allow for defining cell cycle phase from single images

To visualize cell cycle phases in living human ISCs, we adapted the recently published PIP-FUCCI allele as a fluorescent readout for cell cycle phases (Grant, Kedziora et al. 2018). The PIP-FUCCI construct consists of two fluorescent reporters which allow for identification of each cell cycle phase. The Cdt1_30-120_-mVenus fusion has strong fluorescence in G1, rapid degradation (< 30 minutes) at the start of S Phase, then increasing fluorescence in G2 phase through cytokinesis. The Gem_1-110_-mCherry fusion has negligible fluorescence in G1 then fluoresces in the S and G2/M phases through cytokinesis, at which point it rapidly dims (Fig 2A,B). This reporter was chosen for its distinct changes allowing for accurate quantification of all phase lengths using live-cell imaging. As described in (Burclaff, Bliton et al. 2022), we engineered the PIP-FUCCI allele onto a PiggyBac transposase plasmid, transfected this plasmid into human ISCs, and selected stable clones using protocols optimized for primary human ISC monolayers (Breau, Ok et al. 2022). PIP-FUCCI ISCs were thinly seeded onto the collagen hydrogels to allow sufficient space to expand, then imaged starting 24 hours post-plating to mitigate the impact of potential contact inhibition. Since the PIP-FUCCI construct fluoresces distinct color combinations in each cell cycle phase, phases for all cells could be proscribed from a single snapshot (Figure 2C).

**Figure 2:**
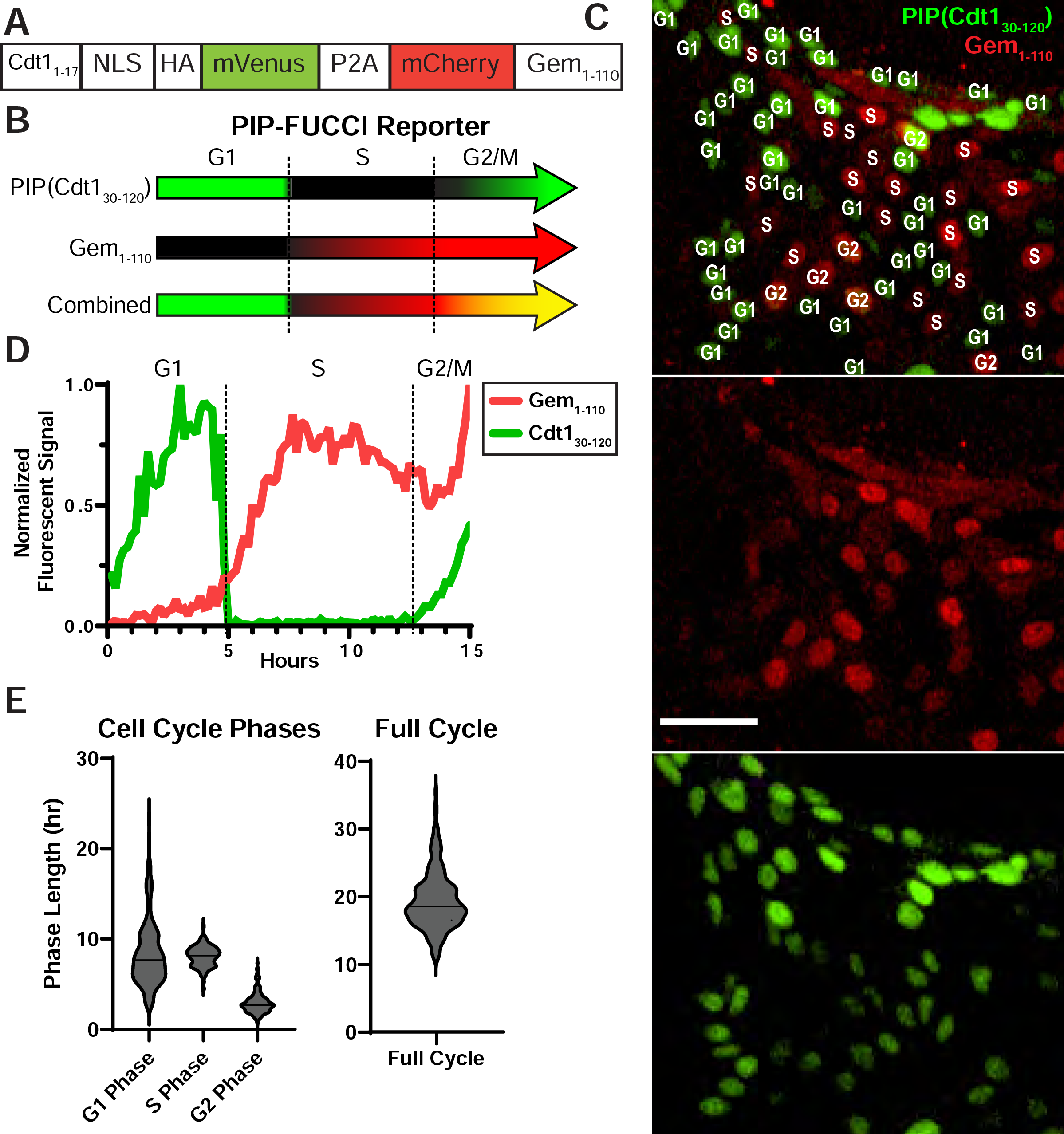
PIP-FUCCI construct and analysis. **A)** Schematic of PIP-FUCCI genetic construct **B)** Schematic of dynamic reporter colors throughout the cell cycle in PIP-FUCCI cells. **C)** Representative fluorescence image of growing PIP-FUCCI human ISC monolayers. Scale bar = 50 μm. The cell cycle phase for each nucleus is superimposed using information from the red and green signal channels. **D)** Example fluorescence data resulting from tracking an individual PIP-FUCCI nucleus over its full cell cycle. **E)** Violin plots denoting lengths of each cell cycle phase across 196 ISCs as quantified using the PIP-FUCCI reporter.

To determine if this platform could be used to directly measure cell cycle phase lengths in freely cycling human ISCs, we live imaged cells for 48 h by confocal microscopy (Supplemental Video). Fluorescence profiles of individual nuclei were tracked across each 10 min frame interval, with changes in mVenus and mCherry levels used to define lengths of each cell cycle phase (Figure 2D). However, as discussed in the original study designing the PIP-FUCCI construct (Grant, Kedziora et al. 2018), a gap exists in early S Phase where both reporters are very dim across 1-3 frames. With no other nucleus marker and given the dynamic movement of primary human ISC monolayers, we were unable to automatically track nuclei through the full cell cycle using publicly available programs. Individual nuclei were tracked by hand through all frames of a cell cycle to observe changes in fluorescent reporters and determine phase lengths (Figure 2D). Our final data analyzing 196 ISCs showed that freely cycling PIP-FUCCI ISCs had a median total cell cycle length of 18.6 hours, with median lengths of cell cycle phases of 7.7 hours for G1 phase, 8.2 hours for S phase, and 2.6 hours for G2/M phases (Figure 2E). Sample files for analyzing cell cycle phases using PIP-FUCCI ISCs are available as Supplementary Files 2-4. Thus, while PIP-FUCCI ISCs can be used to directly quantify cell cycle phase lengths in live imaging, the analysis pipeline is low throughput and requires significant manual input. Instead, this human cell cycle reporter is optimal for defining cell cycle phases of live human ISCs using single snapshot images.

### PIP-H2A human ISC reporters allow for automated cell tracking from live imaging to define cell cycle phases

We next designed a reporter construct to allow for automated analysis of cell cycle phase lengths in live human ISC monolayers while still only using two fluorescent channels. A construct was designed that would label all nuclei with a constitutive fluorescent reporter gene to allow constant tracking of each cell alongside the same PIP-mVenus reporter gene used in the PIP-FUCCI construct to define the precise onset of S and G2 phases. For a bright nuclear reporter, we combined the mScarlet-tagged human histone H2A allele (Bindels, Haarbosch et al. 2017) with the original PIP (Cdt1_30-120_-mVenus fusion), creating a novel PIP-H2A reporter line in our primary human ISCs (Figure 3A). Importantly, this construct is not designed for identifying cell cycle phases using a single snapshot, as cells in both the G1 and G2 phases exhibit the same fluorescence signature (Figure 3B). Rather, the cell cycle reporter gene was designed specifically to track hierarchical cellular division events and quantify cell cycle phase lengths in live imaging. The PIP-H2A cell cycle reporter was optimally designed to define cell cycle phases in live imaged cells: constitutive H2A-mScarlet signal provided the ability to constantly track nuclei and the PIP-mVenus signal enabled precise identification of G1 (high mVenus), S (rapid loss of mVenus), and G2/M phases (increasing mVenus). With the constant expression of H2A-Scarlet, cytokinesis can easily be observed by microscopy to define the beginning and end of full cell cycles. While an additional far-red fluorescent protein could have been added to visually distinguish between G1 and G2, we avoided using a far-red constitutive reporter to leave open a fluorescent channel for additional stains or reporters.

**Figure 3:**
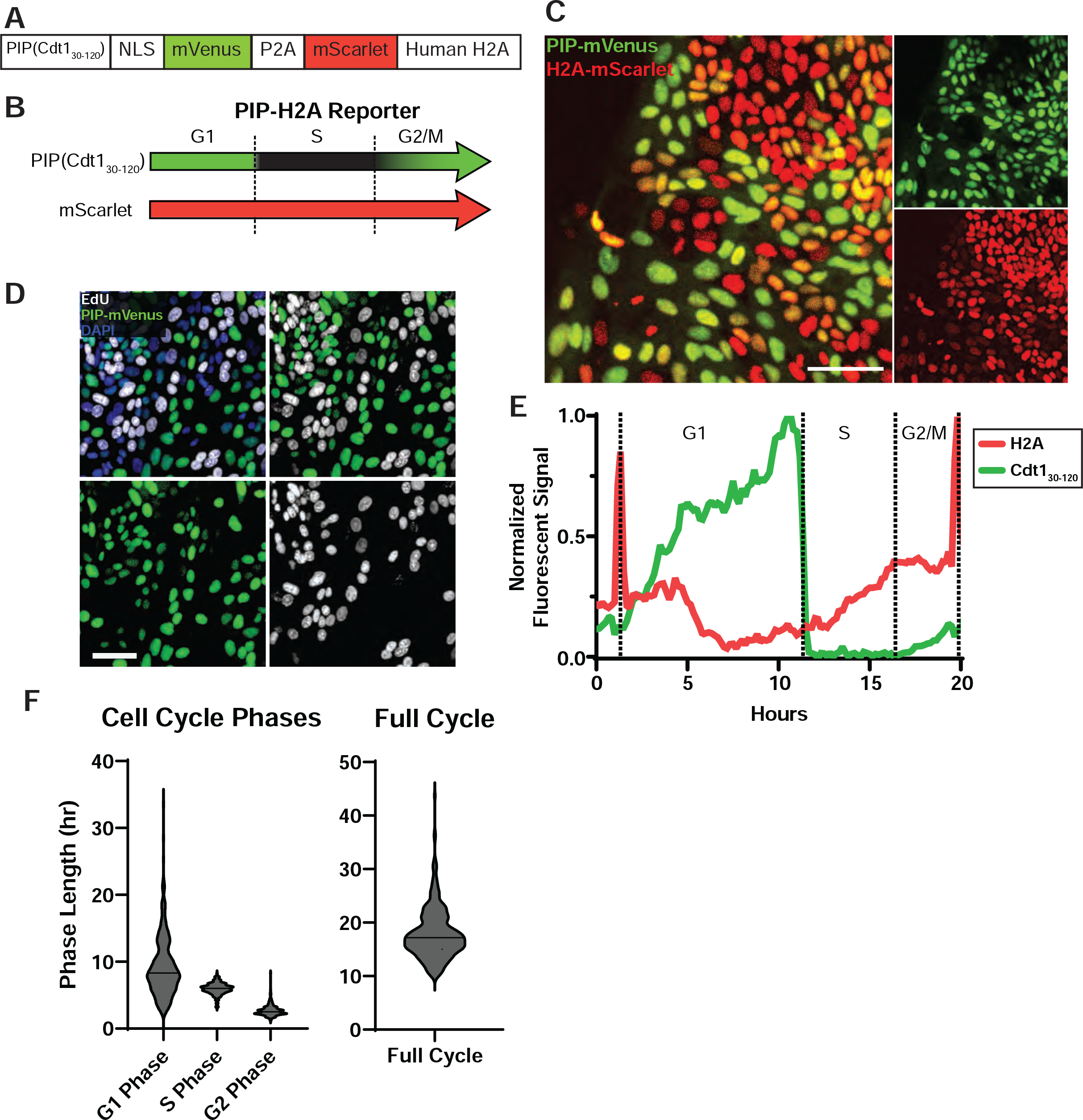
PIP-H2A construct and imaging. **A)** Schematic of the PIP-H2A genetic construct **B)** Schematic of dynamic reporter colors throughout the cell cycle in PIP-H2A cells. **C)** Representative fluorescence image of the PIP-H2A reporter in expanding human ISC monolayers. Scale bar = 50 μm. **D)** PIP-H2A cells stained for EdU following 1 hr EdU pulse. Separate boxes showing individual colors (DAPI, blue; EdU, white; PIP-mVenus, green) display exclusive mVenus or EdU signal in each nucleus. **E)** Example fluorescence data resulting from tracking an individual PIP-H2A nucleus over its full cell cycle. **F)** Violin plots denoting lengths of each cell cycle phase across 284 ISCs as quantified using the PIP-H2A reporter.

Clonal ISCs were generated by electroporation and stable selection of the PIP-H2B construct using established methods (Breau, Ok et al. 2022) followed by functional validation of the reporter construct. The selected clonal cells all showed mScarlet fluorescence, with mVenus fluorescence confined to a subset of cells, as expected (Fig 3C). Interestingly, we noted variable mScarlet signal brightness in these cells that appeared inverse to mVenus signal brightness (Fig 3C,E). To validate that the PIP-mVenus reporter was faithfully marking G1 and G2/M phases but absent in S phase, cells were treated with a one-hour EdU pulse to mark cells in S phase. Fluorescence microscopy confirmed that cells marked by EdU were distinct from cells expressing mVenus (Figure 3D), validating that the construct was accurately reporting the beginning and end of S phase.

To increase the throughput of measuring cell cycle lengths in primary ISCs cells, it was necessary to develop a new automated image analysis pipeline. This approach was designed to identify all nuclei computationally from the live imaging results, track individual nuclei across timepoints, then analyze the green and red signal changes in each nucleus over time. Conceptually, these data for each cell could then be tracked over a full-cell cycle and the time between fluorescent color changes could be aligned to a cell cycle phase. To quantify cell cycle phase lengths in freely cycling primary human ISCs, cells were plated sparsely on pressed collagen chamber slides to avoid contact inhibition, cultured for two days to allow monolayers to establish, then live imaged for 48 hours with imaging at 10-minute intervals. To optimize automatic tracking of nuclei across timeframes, we added the signal intensities from the mScarlet and mVenus channels together. Adding these two nucleus-specific reporter signals resulted in a third channel with strong constant nuclear signal, allowing cells to be tracked computationally throughout the cell cycle and avoiding the manual tracked needed with the PIP-FUCCI reporter.

A software package, CellPose2 (Pachitariu and Stringer 2022), was used to segment nuclei on individual frames of our ISC monolayer live imaging results. Human-in-the-loop training was utilized, with researchers correcting consecutive iterations of the model’s segmentation attempts on these complex images to train the model. Example images are shown in Figure 4A, with the computerized nuclei segmentation shown on the left and the manually corrected image on the right. To track training progression, the initial count of nuclei segmented by the model and the final corrected count were recorded (Figure 4B). In the 5th-7th rounds of training, the model was reaching >95% accuracy, at which point we considered it adequately trained. Once the model was trained using single images, we developed code to analyze all 288 frames of the live imaging file in CellPose2 consecutively (Supplemental File 5), resulting in a TIF file with segmentation across all frames of the live imaging run.

**Figure 4:**
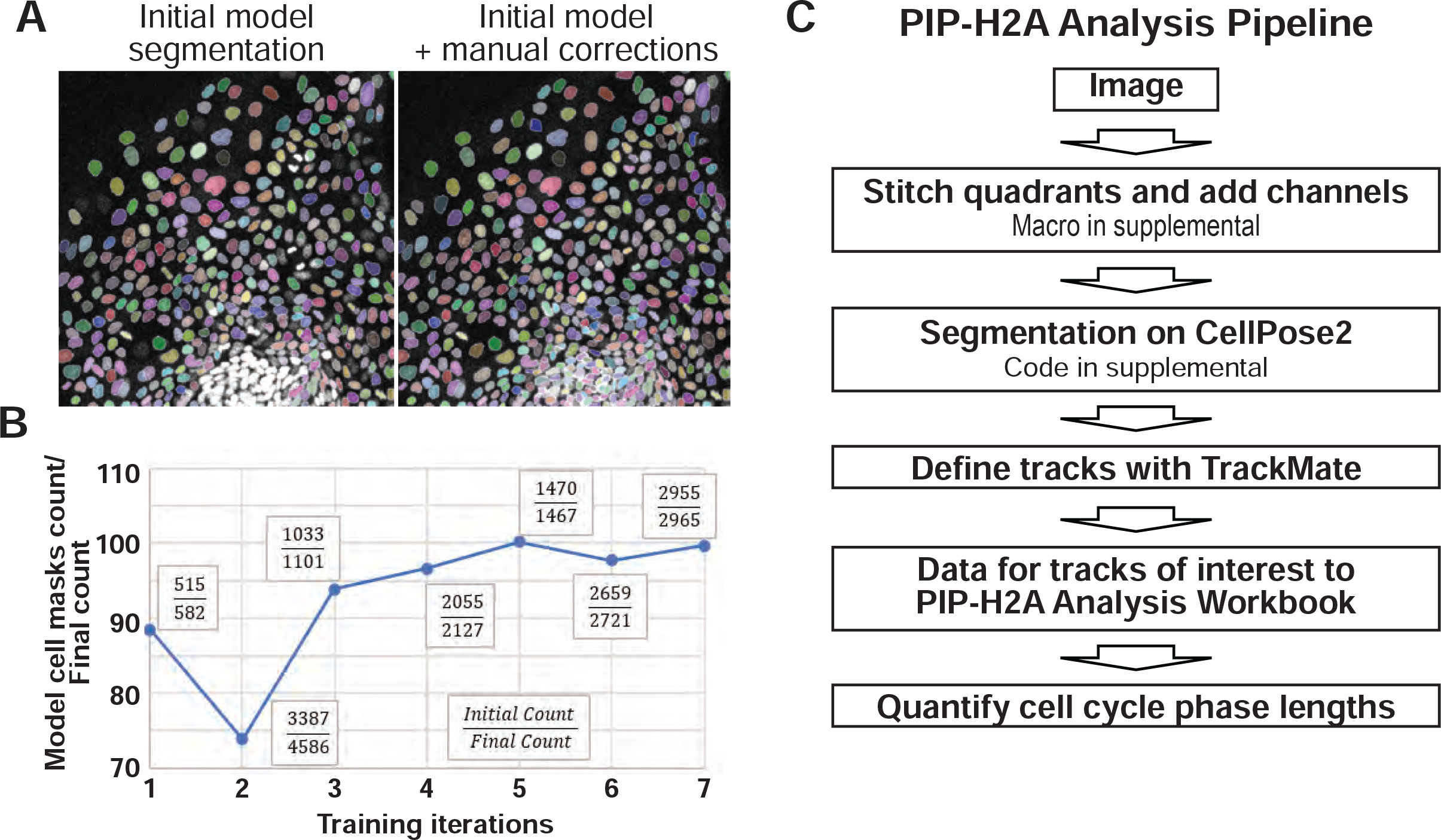
Analysis pipeline for PIP-H2A live imaging. **A)** (Left) Example images showing the attempt by the CellPose2 model to segment nuclei in human ISC monolayers. Segmented nuclei are highlighted with a colored mask; white nuclei lack masks. (Right) Final segmentation following human correction. **B)** Readouts of mask numbers obtained by CellPose2 segmentation modeling/human-corrected final segmentation across 7 iterations of training. **C)** Schematic of the PIP-H2A live imaging analysis pipeline.

Once we obtained segmented files with nucleus masks automatically defined across timepoints, we next needed to analyze reporter signals across time for each nucleus. To do this, we used the TrackMate Fiji plugin (Tinevez, Perry et al. 2017, Ershov, Phan et al. 2022). This program uses the segmentation channel to track individual nuclei across frames based on proximity of each object between frames. To select tracks likely to follow a single nucleus through a full cell cycle, no gaps were allowed between frames and all tracks under 480 minutes were removed from analysis, since our earlier results analyzing the PIP-FUCCI construct found no cell cycle lengths below 500 minutes for all cells analyzed. Tracks under 480 minutes largely represented nuclei tracks that only covered a portion of a cell cycle before being lost due to improper segmentation of the nuclei by CellPose2 or by TrackMate failing to correctly assign the track due to the close proximity and dynamic motion of primary human ISC monolayers, especially around cytokinesis events. To ascertain that all tracks included in our final analysis were of a single nucleus that completed a full cell cycle, tracks for each cell were manually viewed. Our live imaging results showed that the mScarlet-H2A reporter experienced an acute spike in mean intensity as chromatids condensed and split in cytokinesis, allowing mitotic events and total cell cycle length to be readily tracked from the red channel alone (Figure 3E). Our final data analyzing 284 ISCs showed that freely cycling ISCs had a median total cell cycle length of 17.2 hours, with median individual cell cycle phase lengths of 8.3 hours for G1 phase, 6.0 hours for S phase, and 2.5 hours for G2/M phases (Figure 3F). A full pipeline schematic for analyzing PIP-H2A live imaging data is shown in Figure 4C, and sample files are available as Supplementary Files 5-7.

## DISCUSSION

In this study, we genetically engineered primary human ISCs with two cell cycle reporter constructs, developed a microphysiological platform optimized for live imaging these cells, then compiled a pipeline incorporating freely available programs to track cell cycle phase lengths. This platform directly measuring cell cycle phases in freely cycling primary human ISCs represents an important step forward, as ISC cell cycles have only been visualized using model organisms in current literature. Impressively, our human cell cycle data is similar to data reported from intestinal organoids derived from FUCCI2 mice (Matsu-Ura, Dovzhenok et al. 2016). While this study in mice didn’t focus on individual phase lengths, they tracked fluorescent change to report a total cell cycle length of 18.4 h in murine organoids. Not only is this total length very similar to those defined by our human platform, the distributions of the total cycle lengths across ISCs are also similar between studies. They report a large portion of ISCs cycle around 15 h with another notable peak on their histogram arising at slightly over 20 h. This is similar to our data from both PIP-FUCCI (Figure 2E) and PIP-H2A (Figure 3F) human ISCs, with both plots showing a large portion of ISCs cycling about midway between 10 and 20 h and a second population cycling slightly above 20 h. This points to stunning similarities between mouse and human ISCs and demonstrates the strength of tracking cell cycle phases in hundreds of individual ISCs concurrently. Future work may combine our platform with additional staining or lineage reporters to further delineate which cells are proliferating with these two cell cycle lengths.

Several obstacles had to be overcome to engineer systems for analyzing cell cycle phase lengths in primary human ISCs. ISCs are sensitive to platform substrate stiffness, yet microscope objectives often have limited working lengths. We designed a 3D-printed collagen press to make thin, flat patties of soft collagen that support primary ISC growth as monolayers within the observable range. Primary ISC monolayers move dynamically, both as the monolayers spread and migrate and especially when cells or their neighbors undergo cytokinesis. This makes these primary cells more difficult to analyze than many models which use less-motile cells or cells which associate less closely with each other. For accurate automated tracking of these dynamic cells, the nucleus must always be visible fluorescently. As such, we engineered the original PIP-FUCCI cell cycle reporter plasmid to include a nuclear reporter, making the PIP-H2A reporter. Finally, we designed a pipeline combining cutting-edge, yet free and publicly available, computational techniques including CellPose2 and TrackMate to analyze the resulting PIP-H2A ISC live imaging data.

Our PIP-FUCCI and PIP-H2A constructs present many opportunities for use in diverse applications. The PIP-FUCCI construct is optimal for snapshot analyses, where ratios of the cell population in each cell cycle phase can be determined from a single image. Since the analysis can be done in live cells, imaging can be done at different timepoints, for example before and after a drug is added, gene function is induced, or injury occurs. This genetic construct could also be easily scaled up into plates with more wells for drug screens or other large assays, with snapshots across wells analyzed to quickly and easily determine effects on cell cycle phases in primary human ISCs. If direct measurements of cell cycle phase lengths are needed, the PIP-H2A construct is better suited for live imaging. Direct quantifications of cell cycle phase lengths in individual cells can give more detailed information than ratiometric analyses like those done for PIP-FUCCI snapshots or flow cytometry for DNA content, which can be useful when studying agents or proteins expected to regulate cell cycle. Importantly, we avoided the simple strategy of adding a far-red constitutive reporter to the original PIP-FUCCI cells to leave open a fluorescent channel for fluorescent stains or additional reporters to be combined with this powerful reporter in future applications.

While our new automated approach is a significant improvement in resolving human ISCs lengths and represents a substantial increase in throughput compared to the human PIP-FUCCI reporter ISCs, there are some limitations. The main drawback of this system is the movement and proximity of nuclei in the primary human ISC monolayers. Cells migrate quickly, and cytokinesis events occur within the plane of the monolayer, resulting in dynamic movement as nuclei separate and neighboring nuclei are jostled. For these reasons, even after completing seven rounds of training and with CellPose2 accurately defining >95% of nuclei per frame, we watched each analyzed track by eye to ascertain that one single nucleus was followed for a full cell cycle for each track. The dynamics of the H2A-mScarlet signal readout made this easier, as we could quickly scan for tracks that have the tight peaks stereotypical of chromatid formation and separation (Figure 3E). Tracks can also be lost as a result of our stringent filtering rules prohibiting any gaps in the track. We expect these caveats to be overcome as segmentation and tracking technologies improve, and future studies can also use a shorter imaging time (<10 min/frame) to minimize the distances nuclei move between frames.

In all, this study represents a full platform for analyzing cell cycle phase lengths in live primary human ISCs. We provide files for 3D printing the collagen press and for our analysis pipeline to allow other labs to use this technique. Accurately delineating cell cycle phase lengths is important for studying the epithelium in health and disease, identifying cell cycle regulators in ISCs, or defining the effects of drugs and other molecules on cell cycle.

## Methods and Materials

### Tissue procurement, dissection, and crypt isolation

Intestinal crypts were harvested from human intestines following a published protocol (Breau, Ok et al. 2022). Briefly, donor-grade human intestines were received from HonorBridge. The proximal 9 cm of the small intestine as designated duodenum, with jejunum defined as the upper half of the remaining small intestine. A 3x3 cm piece was resected from the middle of the jejunum, then stored in Advanced DMEM/F12 + 10 μM Y27632 and 200 μg/mL Primocin on ice until crypt isolation. The tissue was incubated in PBS + 10 mM N-acetylcysteine for 15 min then transferred to Isolation Buffer (5.6 mM Na2HPO4, 8.0 mM KH2PO4, 96.2 mM NaCl, 1.6 mM KCl, 43.4 mM Sucrose, and 54.9mM d-sorbitol) + 2 mM EDTA + 0.5mM DTT for 30min with gentle rocking at room temperature, followed by vigorous shaking for 2 min. After shaking, the tissue resection was transferred to a new tube of Isolation Buffer + EDTA + DTT and the rocking then shaking were repeated six times, with supernatant for each stored on ice. Following the shakes, supernatants from each round were checked via light microscope for the presence of crypts and villi. Crypt-enriched shakes were pooled, washed in Isolation Buffer, then cultured or frozen

### Tissue Culture

Primary human intestinal epithelial cells were cultured in collagen coated well plates prepared following a published protocol (Wang, DiSalvo et al. 2017, Wang, Gunasekara et al. 2017). Maintenance media was prepared as published in (Hinman, Wang et al. 2021). Briefly, L cells expressing transgenic Wnt3A, Noggin, and R-spondin3 (ATCC CRL-3276) were cultured in Collection Media (20% Tetracycline-negative Fetal Bovine Serum, 1% Glutamax, 1%Pen/Strep, in Adv DMEM/F12) for 12 days, with media collected daily. Maintenance Media consisted of 50% conditioned Collection Media and a final concentration of 2% B-27 Supplement, 5 mM Nicotinamide, 10 mM HEPES, 1 mM Glutamax, 1X Pen/Strep, 0.6125 mM N-Acetylcysteine, 25 μg/mL Primocin, 1.5 μM, 25 ng/mL mEGF, 1 nM Gastrin, and 5 nM Prostoglandin E2. Fresh crypts were plated with 200 mg/mL Primocin, 200 mg/mL Gentamycin, and 0.5 mg/mL amphotericin B for the first week. Cells were grown in a humidified incubator at 37 °C with 5% CO_2_.

### Genetic Engineering and Cloning

All generic engineering was done following established protocols (Breau, Ok et al. 2022). In short, DNA segments of importance were extracted using restriction enzymes or amplified utilizing the CloneAmp HiFi PCR Premix. Plasmids were engineered using an In-Fusion HD Cloning Kit to express the reporter gene of interest between inverted terminal repeats (ITR) to allow for transposing into the genome via super-piggyBac transposase. Plasmids were subsequently harvested from bacterial stocks using the QIAGEN HiSpeed Maxi kit.

The PIP-FUCCI plasmid (Grant, Kedziora et al. 2018) was a gift from Jean Cook and Jeremy Purvis. pPIGA-PHD was a gift from Linzhao Cheng (Addgene plasmid #26778; http://n2t.net/addgene:26778; RRID:Addgene_26778) (Zou, Maeder et al. 2009). sg resistant gamma-tubulin was a gift from Maria Alvarado-Kristensson (Addgene plasmid #104433; http://n2t.net/addgene:104433; RRID:Addgene_104433), and pmScarlet_H2A_C1 was a gift from Dorus Gadella (Addgene plasmid #85051; http://n2t.net/addgene:85051; RRID:Addgene_85051).

Plasmids were introduced into human cells through our optimized electroporation protocol (Breau, Ok et al. 2022), employing the Neon Transfection System with a 100 μL Kit. Cells were suspended in 100 μL of Neon Buffer R, achieving a concentration of 10,000 cells/μL, along with 6 μg of specific plasmid. The Super PiggyBac Transposase Expression Vector, at 5ng/μL, was consistently included in all transfection experiments. The electroporation procedure utilized Neon preset #5 (1,700 V, 1 pulse, 20 ms), after which the cells were promptly transferred to a six-well collagen-coated plate containing 3 mL of Maintenance Media supplemented with Y27632 at a ratio of 1:1000.

After approximately 4-7 days following transfection, colonies emerged and were subjected to selection using Blasticidin (10 μg/mL) for 4-8 days. Surviving colonies were then isolated by digesting the collagen substrate with 100 μL/mL collagenase IV (5,000 U/mL) at 37 °C for 25 minutes. Following this step, the colonies were washed with dPBS, then individual colonies were picked using a 20 μL pipette on a light microscope. These isolated colonies were then placed into separate collagen-coated 48-well plate wells with 300 μL of Maintenance Media and Y27632.

### Staining for proliferation

Stem cell monolayers were pulsed with 10 μM EdU for 1 hour then fixed for 20 min with 4% PFA at room temperature then washed in dPBS and stored at 4 °C until staining. For staining, cells were permeabilized with 0.5% Triton X-100 then stained for EdU Reaction Buffer (4 mM CuSO4, 2 μM Sulfo-CY5-azide, 0.2 M Ascorbic Acid, in PBS) for one hour at room temperature protected from light, washed with PBS, stained with DAPI, then imaged on the Andor Dragonfly Spinning Disk Confocal Microscope. When multiple images are shown in a figure panel, all were taken with the same microscope settings and had the same display adjustments made. Images were analyzed using FIJI software (Schindelin, Arganda-Carreras et al. 2012).

### Printing and Using the Collagen Press

To reproducibly coat an Ibidi μ-Slide 4 Well chamber slide with 0.3 mm collagen, a custom collagen press was created. The press was designed to rest on the walls separating the wells with protruding “feet” that extended into the wells, leaving a 0.3 mm gap for the collagen layer. This press was fabricated using chlorinated polyethylene and an Ultimaker S3 3D printer (Supplemental File 1). The feet of the press were affixed with coverslip glass using Glass Glue, and Rain-X was applied following the manufacturer’s guidelines to allow for removal of the press without pulling up the thin collagen layer from the well. The collagen was prepared according to a published protocol (Wang, DiSalvo et al. 2017, Wang, Gunasekara et al. 2017). Each chamber was supplemented with 250 μL of liquid collagen, the press was immediately positioned on top (as illustrated in Fig 1), then the chamber slides with the press were moved to a 37 °C incubator for 90 min to allow the collagen to solidify. Afterward, the presses were gently detached, and dPBS was added to cover the collagen hydrogel within each chamber. The prepared slides were hermetically stored at room temperature in a sealed zip-top bag until use.

### Live Imaging

Freely-growing monolayers that had been stably transfected with the PIP-H2A fluorescent reporter were seeded onto collagen-coated μ-Slide 4 Well ibiTreat chamber slides for live imaging. Cells were transferred into the pre-prepared chambers 48 hours before imaging, and growth media was changed in the morning prior to the imaging session. Live imaging was performed on an Andor Dragonfly Spinning Disk Confocal Microscope mounted on a Leica DMi8 microscope stand, using a Leica HC PL APO 20x/0.75 LWD air objective. The pinhole size was set to 40 μm. The camera was an Andor iXon Life 888 EM-CCD, with Electron Multiplying Gain set to 150, Horizontal Shift Speed of 10 MHz – 16 bit, 2X Pre Amp Gain, 2.2 μs Vertical Shift Speed, Normal Vertical Clock Voltage and binned 2×2.

To image the mCherry allele in the PIP-FUCCI construct, a 561 nm laser was used for excitation, and light was collected with a 593/43 Semrock emission filter. To image the mScarlet allele in the PIP-H2A construct, a 561 nm laser was used for excitation, and light was collected with a 593/43 Semrock emission filter. For mVenus imaging, common to both constructs, a 514 nm laser was used for excitation, and light was collected with a 538/20 Semrock emission filter. Images had 512 × 512 pixels and the size of each pixel was 1.00 μm. At each position, Z stacks were acquired using a piezo Z stage with 5 μm intervals spanning 75 μm (16 steps). 2×2 montages were acquired at each position. Each Z stack in the 2×2 montage overlapped its neighboring Z stacks by 10%. Two locations with a 2×2 montage Z stack in each of the four wells were acquired per experiment. 2×2 montage Z stacks at each location were acquired every 10 minutes for 48 hours. All images directly compared to each other were acquired using the same settings. Typical settings were 600 ms exposures with 5% laser power for mScarlet, and 200 ms exposures with 1% laser power for mVenus. Temperature was maintained at 37 °C with an Okolab microscope enclosure, with continuous monitoring and feedback. 5% CO2 was warmed to 37 °C in the enclosure and humidified before being delivered to an enclosed stage top holder that contained the sample. To minimize evaporation during the experiment, 8 caps from 15mL falcon tubes were filled with water and placed surrounding the sample inside the stage-top sample holder.

### Live-Imaging Analysis

For the PIP-FUCCI construct: Stitched Z stacks were analyzed using Bitplane Imaris software (version 9.9.1) and Microsoft Excel. A full analysis protocol, a custom Excel analysis workbook, and a sample analyzed Imaris file are included as Supplemental Files 2-4. First, Z slices without nuclear fluorescent signal were cropped out, and all remaining z stacks were max projected along the XY plane. Second, individual nuclei at each timepoint were marked manually using the “spots” function in Imaris. Third, tracks were generated to connect the positions of nuclei marked with spots throughout the experiment. Fourth, mean fluorescent intensities were exported from Imaris into a custom Excel workbook that simplified determination of cell cycle phases. Cells were only tracked from mother cells which divided at least 5 h after the start of imaging to allow cells to acclimate to the imaging environment. Tracking was also only started on mother cell divisions that occurred at least 20 h prior to the end of the video. Cells were only tracked if they stayed in frame and in focus throughout their entire cell cycle, and cells were chosen from a variety of spots across the viewing area and the timelapse for tracking. An average of 90 nuclei were tracked at each location in an experiment (range=59-106 nuclei). The Excel workbook presents graphs of normalized mVenus and mCherry fluorescence. The G1-S transition was defined as the point when mVenus signal dropped over 50%. The S-G2 transition was defined as the point when the mVenus signal began rising above the lowest maintained level again. Final graphs and significance analyses were made using GraphPad Prism 9. Data is shown as violin plots for each cell cycle phase and for total cell cycle duration, with median values shown.

For the PIP-H2A construct: Images were flattened using maximum projection for maximum intensity then stitched using FIJI software, with 10% overlap between quadrants. To maximize nuclear signal for computational tracking, the mScarlet and mVenus channel intensities were added together using FIJI to form a new channel. This summed and stitched channel was imported to CellPose2 for segmentation. An automated Macro to run the previous steps on FIJI is available in Supplemental Figure 5. To train the CellPose2 model, seven individual frames were used, both individual positions and full stitched positions. The “nuclei” option from the model zoo was chosen, with average nuclei diameter set to ‘12’. For each training frame, the model was allowed to predict nuclei segmentation then a researcher would correct the results. Initial and final nuclei counts were recorded for each training session. Upon reaching at least 95% accuracy for three consecutive rounds, the model was considered ready to use. A Jupyter Notebook script was used to run the CellPose2 analysis on all 488 frames of the time-lapse image individually and save the resulting masks as a single TIFF file with 488 frames (Supplemental File 5). Nuclei were tracked across frames using the Fiji Trackmate Plugin. To begin, a four channel TIFF was made combining C1: mScarlet, C2: mVenus, C3: C1+C2, C4: segmentation file from Trackmate for every timepoint. This file was then imported into Trackmate. No cropping was chosen, and the Label Image Detector function was chosen using the segmentation masks in channel 4 to define nuclei. No initial thresholding or filtering was done prior to defining tracks. The “LAP Tracker” was chosen, with specifications set to 10um max distance, Gap close = 7.0, and 0 gaps allowed. After tracks were defined, tracks with total duration <480 min were filtered out as that was shorter than the shortest cell cycle recorded using the PIP-FUCCI reporter. To choose tracks of nuclei which underwent a full cell cycle within the viewing period, Track name, frame, and Max Intensity of C1 values were all exported to Microsoft Excel then filtered for tracks which had at least one point where slope > |300|, following observations of drastically brighter mScarlet signal during cytokinesis (Fig 3 E). Candidate tracks were monitored by eye to ascertain that the track followed a single nucleus through two splitting events, then raw data for the track was copied to an Analysis Workbook in Microsoft Excel (Supplemental File 6-7). The Analysis Workbook presents graphs of normalized mVenus and mScarlet fluorescence. The G1-S transition was defined as the point when mVenus signal dropped over 50%. The S-G2 transition was defined as the point when the mVenus signal began rising above the lowest maintained level again. M/C1 transition was defined by the acute peaks in mScarlet intensity. Final graphs and significance analyses were made using GraphPad Prism 9. Data is shown as violin plots for each cell cycle phase and for total cell cycle duration, with median values shown.

## Supporting information

Supplemental File 1: Collagen press design file

Supplemental File 2: PIP-FUCCI analysis protocol

Supplemental File 3: PIP-FUCCI empty analysis workbook

Supplemental File 4: Example PIP-FUCCI analysis workbook

Supplemental File 5: Code files for PIP-H2A live imaging analysis

Supplemental File 6: PIP-H2A empty analysis workbook

Supplemental File 7: Example PIP-H2A analysis workbook

Supplemental Table: Reagents and Materials

Supplemental Video: Live imaging of cycling PIP-FUCCI cells

## ACKNOWLEDGEMENTS

The authors first and foremost thank the human donor and their family for the gift of tissue. The authors thank Steven Emanual through the Biomedical Engineering Department at UNC Chapel Hill for help with 3D printing the collagen presses. We thank Jean Cook and Jeremy Purvis for supplying the PIP-FUCCI construct and help with analyzing cell cycle. The Microscopy Services Laboratory, Department of Pathology and Laboratory Medicine, is supported in part by the P30 CA016086 Cancer Center Core Support Grant to the UNC Lineberger Comprehensive Cancer Center. The Andor Dragonfly microscope was funded with support from National Institutes of Health grant S10OD030223.

This research was supported by a CGIBD pilot grant through funding from the National Institutes of Health, P30 DK034987, T32DK07737, and F32DK124929 (J.B.), T32GM133364 (K.A.B.), F30DK126307 (M.T.O.), R01DK115806 and R01DK109559 (S.T.M.), F30AI172230 (C.Y.T.), and the Katherine E. Bullard Charitable Trust for Gastrointestinal Stem Cell and Regenerative Research.

