## Supplemental File 2: PIP-FUCCI analysis protocol for "An in vitro platform for quantifying cell cycle phase lengths in primary human intestinal stem cells"

### PIPFUCCI Analysis workflow

#### Part 1 - Imaris workflow

1- **Max project the XY plane along the Z axis, using Imaris (Image Processing/Project to 2D).**
   a. Consider cropping out Z planes that do not have nuclei for the entire experiment before this step.
   b. Max projection simplifies the data from 3D to 2D. Because cells do not form piles, this does not introduce problems in this dataset.

**2- Mark nuclei using the spots tool (one daughter pair at a time)**
   a. Create a spots object. Click on “Skip automatic creation, edit manually”.
   b. Make sure you are on the edit tab (pencil).
   c. Set the default diameter of the spots. To do this, click on the top right pointer tool. Using shift and the scroll wheel will change the size of a bounding box which will in turn determine the size of the spots. Notes:
      i. Once a spot is placed, the diameter will show up in the edit tab.
      ii. The size of the spot will determine the area over which the intensity in a nucleus is averaged. Too small will make it very noisy. Too big might overlap with other nuclei and contaminate the signal. Nuclei vary in size between 5-15 um. A reasonable size would be 5.15um (not every size is available given the fixed pixel size)
      iii. To avoid changing the diameter by mistake halfway through the track, switch to the second pointer tool in the top right (the one with the circle). With this tool, shift-scrolling changes the size of the selection box, NOT the actual spots.
   d. Before adding spots, check that they are trackable through the entire cell cycle. Look for:
      i. Red blob that separates into two green blobs
      ii. Collapse in green signal
      iii. Rise in red signal (green signal also rises a bit at the end, marking start of G2)
      iv. Red blob splits again
   e. Add spots by shift-left clicking while in the edit tab. Add spots in pairs of sister cells that have just appeared from division of a mother cell. If the fluorescence disappears, move forward and try to infer where the spot might have been in between.
   f. Make sure the size of the selection circle is such that it does not intersect with other spots or tracks. If it does, you will not be able to add a spot. You can change the size of the selection circle to avoid this and/or turn off the display of tracks in the spots setting tab (Tracks Display off)

**3- Track the spots (one daughter pair at a time)**
   a. Highly recommended to track daughter pairs one at a time, to make sure there are no gaps in the tracking.
   b. To track, go to the creation tab in the spots object. Click on Recompute, tracks.
   c. Enter the following parameters:
      i. Algorithm: Brownian motion (this means that the tracking algorithm does not assume the spots move in a predictable way).

ii. Max distance: 8um works well. This is the maximum distance a spot can move between frames and still be connected. Too small means tracks are fragmented. Too large can cause tracks to get confused.
iii. Max Gap Size: 0. This means every frame has a spot.
iv. Fill gaps with all detected objects: UNCHECKED

d. Click on the blue right arrow.
e. Notes:
    i. This will recalculate ALL tracks in the spot object. Ergo, multiple tracks on multiple nuclei are recalculated.
    ii. This can scramble track ID numbers. Until all tracks are calculated for a given spots object, do not do anything with those IDs in subsequent analysis.
f. Quality control for tracks:
    i. In the settings tab of the spots object use
        1. Whichever tracks display option you prefer: full tracks or dragon tails (and for how long).
        2. The tracks displacement display arrows can also be useful
        3. Points style center point can also be useful; reducing the size of the sphere can allow better visibility.
    ii. Check that the filtering gives the expected number of spots. For example, if you have 5 pairs of cells you should have 10 tracks. If not, adjust the filtering.
    iii. Move the time slider to check whether the tracks are all continuous (no gaps) and nothing got crossed.
    iv. If there are gaps:
        1. Check that there is a spot in each frame.
        2. Consider rerunning with a larger max distance parameter.
    v. If tracks get "crossed" between spots:
        1. Consider rerunning with a smaller max distance parameter.
        2. If tracks cross and it is hard to tell, consider dropping the spot or editing the track manually.
g. Click on the orange circle with an X: "Cancel: keep the created objects and terminate the wizard". This keeps the tracks you just made.

**4- If everything looks good SAVE.**

**5-** IMPORTANT: each nucleus has a unique TrackID identifier. These can be used to unambiguously associate sister, mother and daughter cells outside of Imaris. There is also an ID for each spot, but these change for each frame.

**6- Export the data:**

a. Select the spot object whose tracks you want to export.
b. Go to the filter tab of the spot object.
c. Add a filter (it does not matter what it is).
d. Adjust the filter so that all objects are selected. This step is critical for exporting everything. If you don't do this, you will only be able to export data in the desired format from spots visible at a given timepoint. Since not every spot is on each timepoint, this would make exporting very cumbersome.

e. Go to the statistics tab of the spot object. Select Specific Values, Intensity Mean Ch=1 Img=1 (this is the Red channel).
f. Click on the bottom button “Export data for plotting”. Use xls format. Add Ch1 to the name
g. Select Specific Values, Intensity Mean Ch=2 Img=1 (this is the green channel).
h. Click on the bottom button “Export data for plotting”. Use xls format. Add Ch2 to the name
i. Quality control: Your Excel spreadsheets should have a number of data columns equivalent to the total number of spots in the spots object. Example: 5 pairs of daughter cells should be 10 columns.

#### Part 2 - Excel workflow

1- **Copy/paste the data correctly into an empty Excel analysis workbook**
   a. Workbook to use is “Supplemental File 3 - Empty Analysis Workbook for PIPFUCCI”
   b. Note that timepoint 1 may not be time=0 if there was no spot detected on the very first frame of the experiment; the workbook will adjust as needed.
   c. Copy all columns from the worksheets Imaris generated and paste into cell B1 of the worksheets “Intensity Mean Ch=1 Img=1” and “Intensity Mean Ch=2 Img=1”.
   d. Once all the data is pasted, “save as” with a different name.

2- **Basic Excel workbook navigation**
   a. “Intensity Mean Ch=1 Img=1” and “Intensity Mean Ch=2 Img=1” worksheets contain the raw data, plus a reference number in the first column. Beyond copy-pasting the data here, there is no need to interact with these sheets.
   b. “Worksheet” is where all the calculation and lookup is done. Do not modify anything here.
   c. “Analysis” is where the user can explore data from each cell pair, input lineage relationships, input transition points in the cell cycle, and extract basic statistics. <u>Gray cells are where the user can or must input data.</u>

**3- Cell cycle inputs in the analysis worksheet**
   a. Input desired cell pair. It strongly recommended to go in order, from 1 to the total number of cell pairs marked in Imaris.
   b. If the cell pair number has data for it, 4 graphs show with the normalized green and red signal vs total experiment time and vs relative experiment time. The latter graphs are more useful for determining when the G1->S and S->G2 transition points occur.
   c. Input the timepoint where green signal goes down (G1->S), and when it starts going back up (S->G2). If you want to be precise (respecting the 10 min timing of the experiment), use numbers ending in 0, 0.17, 0.33, 0.5, 0.67, and 0.83.
   d. If you don’t input a green signal rise the spreadsheet assumes you could not determine the S->G2 transition. Hence S, G2, and total cell cycle duration are not knowable.

**4- Lineage inputs in the analysis worksheet**
   a. If you have lineage relationships between tracks you can input them as daughter of, mother of 1, mother of 2.

**5- Extracting cell cycle statistics across multiple cells**
   a. Once you’ve completed all inputs for each cell pair, copy the light orange cells and paste in the gray area on the bottom, <u>as values.</u>

b. As you paste values from each cell pair, box plots of G1, S, G2, total cell cycle and cell cycle vs start time in experiment are populated.

**6- Validation, error checking and limitations**

a. Once you input green signal drop and rise times, cell cycle parameters are calculated automatically. If values are outside of 2 standard deviations from the mean, an alert will show up. If values are normally distributed you should expect around 4% (one out of 20ish) of them to fall outside of 2 standard deviations, but if you see this alert it can be a clue that there was a typo.
b. Once you have completed all cell pairs, if you want to go back to one you've already filled out, the grayed cells will be incorrect. Adjacent cells will show the correct values if the cells have already been analyzed.
c. The worksheet assumes 288 frames spaced 10 minutes apart.
d. The worksheet can accommodate approximately 500 cell pairs.
e. It is recommended to use one worksheet per Imaris image analyzed. Data can be collated between repetitions and groups in subsequent analysis spreadsheets.
f. Spreadsheet was designed for use on a 27-inch monitor and is best viewed in that format.
