## Supplemental File 5: Code files for PIP-H2A live imaging analysis for "An in vitro platform for quantifying cell cycle phase lengths in primary human intestinal stem cells": Cellpose segmentation of timelapse TIFF.ipynb

```
# %% [markdown]
# Cellpose segmentation

# %% [markdown]
Notebook showing how to segment cells using Cellpose.

# %% [code]
import os
import sys
import numpy as np
from tifffile import imread, imwrite
from cellpose import models

# %% [code]
import torch
torch.cuda.get_device_name(0)

# %% [code]
# load cellpose model
model_path = r'D:\User data\Joseph Burclaff\models\CP_20230707_090400'
model = models.CellposeModel(gpu=True, pretrained_model = model_path)

# %% [code]
im_path = r'D:\User data\Joseph Burclaff\COMPILED TRACKMATE FOR COTTON 2023\MJC005 full scan_2023-08-18_09.53.45_Field_8\MaxProjectSumONLYStitch\MJC005 full scan_2023-08-18_09.53.45_F28to31.tif'
save_dir = r'D:\User data\Joseph Burclaff\COMPILED TRACKMATE FOR COTTON 2023\MJC005 full scan_2023-08-18_09.53.45_Field_8\MaxProjectSumONLYStitch'
masks_name = 'MJC005_Pos4_Segmented.tif'

# %% [code]
im = imread(im_path)

# %% [code]
im.shape

# %% [code]
# loop for segmentation 

labels_list = []
for im_frame in im : 

    # segment the right plane
    labels, _, _ = model.eval(im_frame, diameter=14, channels=[0,0])
    
    labels_list.append(labels)

# %% [code]
# save segmentation

masks = np.array(labels_list)

save_path = os.path.join(save_dir,masks_name)
imwrite(save_path,masks.astype('uint16')) 

# %% [code]


# %% [code]
```
