## Supplemental File 5: Code files for PIP-H2A live imaging analysis for "An in vitro platform for quantifying cell cycle phase lengths in primary human intestinal stem cells": Macro for stitching imaging quadrants.ijm

```
//Ask for directory with files
Dialog.create("Important note on data for this macro");
Dialog.addMessage("\nYou will be asked to select a directory with non-stitched Imaris files.\n \n* That Directory must ONLY contain the non-stitched Imaris files and no other files\n \n* Works with up to 99 positions");
Dialog.show();
inputDir = getDirectory("Select directory which non-stitched Imaris files");

//Get list of files in input directory
inputlist = getFileList(inputDir);

//Create new directories in which to place output
File.makeDirectory(inputDir+"MaxProject");
outputDirMax = inputDir +"MaxProject"+File.separator;
File.makeDirectory(inputDir+"MaxProject3C");
outputDirMax3C = inputDir +"MaxProject3C"+File.separator;
File.makeDirectory(inputDir+"MaxProject3CStitch");
outputDirMax3CStitch = inputDir +"MaxProject3CStitch"+File.separator;
File.makeDirectory(inputDir+"MaxProjectSumONLYStitch");
outputDirMaxSUMonlyStitch = inputDir +"MaxProjectSumONLYStitch"+File.separator;

//Ask for Macro parameters
Dialog.create("Macro Parameters");
Dialog.addNumber("Time Interval (min):",10);
Dialog.addNumber("Pixel size (um):",0.629);
Dialog.addNumber("Number for first tile in mosaic (no leading zeros; example: F04->4)",0);
Dialog.addNumber("Number of columns in mosaic",2);
Dialog.addNumber("Number of rows in mosaic",2);
Dialog.addNumber("Tile overlap (%)",5);
Dialog.show();

TimeInterval = Dialog.getNumber();
PixelSize = Dialog.getNumber();
FirstTile = Dialog.getNumber();
MosaicColumns = Dialog.getNumber();
MosaicRows = Dialog.getNumber();
TileOverlap = Dialog.getNumber();

//Start timer
starttime = getTime();

//Prepare to run all functions
close("*");
setBatchMode(true); 

//Run the MaxProject function on all files in the directory

print("Making Max Projections");

pretimeallmax = getTime();

for (i = 0; i < inputlist.length; i++){
			print("MaxProjecting Image # "+(i+1));
			pretimemax = getTime();
	        MaxProject(inputDir, outputDirMax,inputlist[i],PixelSize);
	        posttimemax = getTime();
	        elapsed = posttimemax - pretimemax;
			print(elapsed/60000+" min to max project image");
}

posttimeallmax = getTime();
elapsedtimeallmax = posttimeallmax - pretimeallmax;
print(elapsedtimeallmax/60000+" min to max project all images");

//Get list of files in maxProject directory
outputMaxlist = getFileList(outputDirMax);

//Run the C1+C2 function on all files in the Max projected directory

print("Adding C1 and C2");

pretimeallsum = getTime();

for (i = 0; i < outputMaxlist.length; i++){
			print("Adding Image # "+(i+1));
			pretimesum = getTime();
	        sumC1C2(outputDirMax, outputDirMax3C,outputMaxlist[i]);
	        posttimesum = getTime();
	        elapsed = posttimesum - pretimesum;
			print(elapsed/60000+" min to sum C1 and C2");
}

posttimeallsum = getTime();
elapsedtimeallsum = posttimeallsum - pretimeallsum;
print(elapsedtimeallsum/60000+" min to sum C1 and C2 in all images");

//Run the stitching function on all files in the Max projected directory

print("Stitching");

pretimestitch = getTime();

stitcher(outputDirMax3C,outputDirMax3CStitch,outputDirMaxSUMonlyStitch,FirstTile,MosaicColumns,MosaicRows,TileOverlap,PixelSize,TimeInterval);

posttimestitch = getTime();
elapsedtimestitch = posttimestitch - pretimestitch;
print(elapsedtimestitch/60000+" min to stitch");

setBatchMode(false);

currenttime = getTime();
elapsedtimeall = currenttime - starttime;
print(elapsedtimeall/60000+" min total processing time");
print(elapsedtimeallmax/60000+" min to max project all images :"+elapsedtimeallmax/elapsedtimeall*100+" % of total");
print(elapsedtimeallsum/60000+" min to sum C1 and C2 in all images :"+elapsedtimeallsum/elapsedtimeall*100+" % of total");
print(elapsedtimestitch/60000+" min to stitch :"+elapsedtimestitch/elapsedtimeall*100+" % of total");

// Definition of functions

// Definition of function that does the max projection

function MaxProject(inputDir, outputDir,filename,PixelSize) {
	close("*");
	//import file
	run("Bio-Formats Importer", "open=["+inputDir+filename+"]");
	run("Z Project...", "projection=[Max Intensity] all");
	//Adjust properties of image with proper scaling
	getDimensions(w, h, channels, slices, frames);
	nchannels = channels;
	nframes = frames;
	run("Properties...", "channels="+nchannels+" slices=1 frames="+nframes+" pixel_width="+PixelSize+" pixel_height="+PixelSize+" voxel_depth="+PixelSize+" frame=["+TimeInterval+" min]");
	//create name for file saving
	savename = File.getNameWithoutExtension(inputDir + filename);
	//save max projection
	print("Saving Max Projection");
	saveAs("Tiff", outputDir + savename);
	close("*");
}

// Definition of function that does the C1+C2

function sumC1C2(inputDir, outputDir,filename) {
	close("*");
	//open file
	open(inputDir+filename);
	//create name for tracking channel files
	workname = File.getName(inputDir + filename);
	//convert to 32-bit
	run("32-bit");
	//split channels
	run("Split Channels");
	//Add C1 and C2 in 32 bit
	imageCalculator("Add create 32-bit stack", "C1-"+workname,"C2-"+workname);
	//Create new 3C stack
	run("Merge Channels...", "c1=[C1-"+workname+"] c2=[C2-"+workname+"] c3=[Result of C1-"+workname+"] create keep");
	//switch display colors to magenta (C1, RFP), yellow (C2, GFP) and cyan (C1+C2) and do display autocontrast
	Stack.setChannel(1);
	run("Magenta");
	run("Enhance Contrast", "saturated=0.35");
	Stack.setChannel(2);
	run("Yellow");
	run("Enhance Contrast", "saturated=0.35");
	Stack.setChannel(3);
	run("Cyan");
	run("Enhance Contrast", "saturated=0.35");
	//save 3C stack
	print("Saving 3C stack");
	saveAs("Tiff", outputDir +workname);
	close("*");
}

// Definition of function that does the stitching

function stitcher(inputDir, outputDir3C, outputDir1C, first, columns, rows, overlap, pixel, time) {
	//create name for saving
	inputlist = getFileList(inputDir); //create list of all files in input directory
	filename=inputlist[0]; //extract name of first file
	filename=substring(filename,0,lengthOf(filename)-6); //extract name of first file up to the F marking the position
	savename=filename+first+"to"+(first+columns*rows-1)+".tif"; //give stitched file an informative name
	
	//Windows backlash issue: swap backslashes in path with forward slashes so that macro can work
	fixPath = replace(inputDir, "\\", "/");
	
	//stich
	run("Grid/Collection stitching", "type=[Grid: snake by columns] order=[Up & Right] grid_size_x="+columns+" grid_size_y="+rows+"  tile_overlap="+overlap+"  first_file_index_i="+first+"  directory=["+fixPath+"] file_names=["+filename+"{ii}.tif] output_textfile_name=TileConfiguration.txt fusion_method=[Linear Blending] regression_threshold=0.30 max/avg_displacement_threshold=2.50 absolute_displacement_threshold=3.50 computation_parameters=[Save computation time (but use more RAM)] image_output=[Fuse and display]");
	
	//OPTIONAL reorder hyperstack, if stitching plugin screws this up. If so, uncomment next two lines.
	//run("Re-order Hyperstack ...", "channels=[Channels (c)] slices=[Frames (t)] frames=[Slices (z)]");
	//switch display colors to magenta (C1, RFP), yellow (C2, GFP) and cyan (C1+C2) and do display autocontrast
	
	//Convert to 16 bit, with appropriate scaling
	//Turn off scaling for conversion from 32-bit to 16-bit to avoid rescaling of raw data in C1 and C2
	run("Conversions...", " ");
	//split channels to convert one at a time, not scaling C1 and C2, but scaling C3:C1+C2
	run("Split Channels");
	selectWindow("C1-Fused");
	run("16-bit");
	selectWindow("C2-Fused");
	run("16-bit");
	selectWindow("C3-Fused");
	//Check whether any pixel intensities are above the 16 bit limit (65535)
	//If so, rescale when converting from 32 bit to 16 bit using an appropriate conversion factor, and inform the user
	//If not, do not rescale the 32 bit sum and do a direct conversion to 16 bit,
	Stack.getStatistics(voxelCount, mean, min, max, stdDev);
	print("Max pixel value of sum C1 and C2 was "+max);
	conversionfactor = 65535/max;
	if(max>=65535){
		run("Multiply...", "value="+conversionfactor+" stack");
		print("Sum of C1 and C2 was rescaled down to 16-bit scale with a 65535/max ("+conversionfactor+") conversion factor"); //informs user that an image was scaled	
	};
	//run("16-bit"); //converts 32 bit to 16 bit
	run("16-bit");
	run("Merge Channels...", "c1=C1-Fused c2=C2-Fused c3=C3-Fused create");

	//Display adjustments
	Stack.setChannel(1);
	run("Magenta");
	run("Enhance Contrast", "saturated=0.35");
	Stack.setChannel(2);
	run("Yellow");
	run("Enhance Contrast", "saturated=0.35");
	Stack.setChannel(3);
	run("Cyan");
	run("Enhance Contrast", "saturated=0.35");
	
	//Adjust properties of image with proper scaling
	getDimensions(w, h, channels, slices, frames);
	nchannels = channels;
	nframes = frames;
	run("Properties...", "channels="+nchannels+" slices=1 frames="+nframes+" pixel_width="+pixel+" pixel_height="+pixel+" voxel_depth="+pixel+" frame=["+time+" min]");
	Stack.setXUnit("micron");
	
	//save 3C stitched stack
	print("Saving 3C stitch");
	saveAs("Tiff", outputDir3C + savename);
	//save 1C (sum of C1 and C2) stitched stack
	run("Make Substack...", "channels="+nchannels+" frames=1-"+nframes+"");
	print("Saving summed channel stitch");
	saveAs("Tiff", outputDir1C + savename);
	close("*");
}
```
