## Supplemental File 5: Code files for PIP-H2A live imaging analysis for "An in vitro platform for quantifying cell cycle phase lengths in primary human intestinal stem cells": readme - PIP-H2A image analysis pipeline.txt

Image analysis pipeline for PIP-H2A platform

Timelapse images were otained as 2x2 grids. 
1) Put the non-stitched quadrants into a folder with no other files present.

2) To stitch the quadrants into a single image with 10% overlap and add C1+C2 to create C3 for optimal segmentation, 
use the Macro provided in this folder named "Macro for stitching imaging quadrants.ijm" with FIJI.

3) To segment the nuclei from the resulting files, use the included python code named "Cellpose segmentation of timelapse TIFF.ipynb".

4) The included code text includes file locations from our directory, replace with the location of your files and a desired destination.
4a) For segmentation, use the output file from (2) labeled "MaxProjectSumONLYStitch".
4b) Our trained CellPose2 model is included in the zip file named "CP_20230707_090400".

5) Run resulting file on CellPose2 using a trained model with cell diameter = 12.

6) Use FIJI to combine resulting segementation file with the 3-channel TIF file made in (2) to create a 4-channel file with segmentation as C4.

7) Open 4-Channel TIFF in FIJI, select Trackmate PlugIn.
7a) Select Label Image Detector, use Channel 4 for masks with simplify controus checked, use simple LAP tracker.
7b) We used max distance = 10um, 0 Gaps allowed, then filtered out tracks under 480 min.
7c) To visualize individual tracks, in Dispaly Tracks, select Show Selection only, with Fade range at 30 time points.
7d) Select "Tracks of Selection" option before viewing tracks to avoid stystem crashing.

8) To optain raw data to analyze tracks, choose "spots" then export the window to .csv.
8a) Mean Intenisty for C1 (mScarlet-H2A) can be graphed to visually select for tracks with tight peaks indicating cell splitting events.  
8b) We recommend watching each track of interest by eye to ascertain that the mask didn't switch between nuclei.

9) To analyze cell cycle phases using our example workbook, copy all data for track of interest, past all into gray columns of the workbook (starting at column AP).
9a) Graphs will autopopulate, fill in cell cycle start/end (red peaks), G1->S (green decline), and S->G2 (green increase) by hand.
